# Hedgehog signaling regulates neurogenesis in the larval and adult zebrafish hypothalamus

**DOI:** 10.1101/740613

**Authors:** Ira Male, A. Tuba Ozacar, Rita R. Fagan, Matthew Loring, Meng-Chieh Shen, Veronica Pace, Chris Devine, Alyssa Lutservitz, Rolf O. Karlstrom

## Abstract

While neurogenesis in the adult hypothalamus is now known to be essential for proper function, the cell-cell signaling events that regulate neurogenesis in this evolutionarily conserved brain region remain poorly understood. Here we show that Hedgehog (Hh)/Gli signaling positively regulates hypothalamic neurogenesis in both larval and adult zebrafish and is necessary and sufficient for normal hypothalamic proliferation rates. Hedgehog-responsive cells are relatively rapidly proliferating pluripotent neural precursors that give rise to dopaminergic, serotonergic, and GABAergic neurons. *in situ* and transgenic reporter analyses revealed substantial heterogeneity in cell-cell signaling within the hypothalamic niche, with slow cycling Nestin-expressing cells residing among distinct and overlapping populations of Sonic Hh (Shh)-expressing, Hh-responsive, Notch-responsive, and Wnt-responsive radial glia. This work shows for the first time that Hh/Gli-signaling is a key component of the complex cell-cell signaling environment that regulates hypothalamic neurogenesis throughout life.

## Introduction

The hypothalamus, located in the ventral-most region of the diencephalon, is among the most ancient and evolutionarily conserved parts of the vertebrate brain (Nieuwenhuys, Donkelaar, Nicholson, & SpringerLink (Online service), 1998). This brain region regulates metabolism, circadian rhythms, autonomic function, and a wide range of behaviors linked to survival including feeding, sleep-wake cycles, and reproduction (Saper & Lowell, 2014). The endocrine hypothalamus secretes neurohormones that regulate the endocrine output of the anterior pituitary gland (Clarke, 2015) while also modulating brain activity through the production of a variety of neuromodulators (e.g. Giardino & de Lecea, 2014). Because of this critical role in basic metabolic processes and behaviors, dysfunction of the hypothalamus is associated with a wide range of human disease, including reproductive impairments, obesity and feeding disorders, abnormal energy metabolism, diabetes insipidus, and sleep disorders, among others (Fliers, 2014; Saper & Lowell, 2014). The hypothalamus is also affected in many neurodegenerative disorders, and alterations in adult neurogenesis in this brain region are likely linked to pathophysiologies accompanying these diseases (Ishii & Iadecola, 2015; Vercruysse, Vieau, Blum, Petersen, & Dupuis, 2018; Winner & Winkler, 2015).

Life-long hypothalamic neurogenesis has now been documented in rodents (Ming & Song, 2011; Yoo & Blackshaw, 2018), sheep (Migaud et al., 2010), zebrafish (Schmidt, Strahle, & Scholpp, 2013; X. Wang et al., 2012), and likely humans (Pellegrino et al., 2018). Additionally, there is growing evidence that hypothalamic neurogenesis plays a role in growth and tissue maintenance as well as hypothalamic function (Kokoeva, Yin, & Flier, 2005; Lee & Blackshaw, 2012; Migaud, Batailler, Pillon, Franceschini, & Malpaux, 2011; Xie & Dorsky, 2017). As with other adult neurogenic zones, hypothalamic neurogenesis occurs in the ventricular regions, namely the posterior recess (PR) and lateral recess (LR) of the hypothalamic (3^rd^) ventricle. Similar to more dorsal neurogenic zones, hypothalamic proliferation requires highly coordinated regulation of cell proliferation and differentiation within a discrete population of progenitors. Cell-cell signaling systems known to regulate nervous system induction and patterning during embryogenesis continue to play a key role in controlling this adult stem cell proliferation and differentiation in other regions of the brain. These include Notch, Fibroblast Growth Factor (FGF), Wnt, Hedgehog (Hh), and Bone Morphogenetic (BMP) signaling systems, among others (Anand & Mondal, 2017; Kizil, Kaslin, Kroehne, & Brand, 2012; Obernier & Alvarez-Buylla, 2019; Petrova & Joyner, 2014). Heterogeneity in both the neural stem cell populations and in cell-cell signaling systems helps control the range of differentiated cell types that are produced in stem cell niches (Ceci, Mariano, & Romano, 2018; Chaker, Codega, & Doetsch, 2016; Lim & Alvarez-Buylla, 2016; Marz et al., 2010). Determining how this heterogeneity contributes to brain growth and adult neurogenesis remains a major challenge in the field.

The Hedgehog (Hh)/Gli signaling system has been extensively characterized for its role in regulating cell proliferation, differentiation, and survival during embryogenesis (Briscoe & Therond, 2013; Varjosalo & Taipale, 2008). Secreted proteins of the Hh family also act through the Gli transcription factors to regulate neural stem cell proliferation in the ventricular-subventricular zone of the hippocampus (Alvarez-Buylla & Ihrie, 2014; Daynac et al., 2016; Lai, Kaspar, Gage, & Schaffer, 2003; Palma et al., 2005; Petrova & Joyner, 2014). Misregulation of Hh signaling is linked to neural tumors including glioblastoma and medulloblastoma (Raleigh & Reiter, 2019; Wechsler-Reya & Scott, 2001) and has been implicated in mediating Parkinson’s disease symptoms and restoring nigrostriatal dopaminergic neurons (Gonzalez-Reyes et al., 2012), but if and how Hh signaling regulates neurogenesis in the adult hypothalamus is not well understood. Given the central role of the hypothalamus in regulating metabolism and other basic physiological processes, understanding the role of Hh/Gli signaling in hypothalamic neurogenesis and its relation to growth, tissue maintenance and plasticity could inform our understanding of a broad range of metabolic diseases associated with this brain region.

The zebrafish brain, which maintains up to 16 proliferative zones throughout life, has become an important model for studying adult neurogenesis (Anand & Mondal, 2017; Grandel, Kaslin, Ganz, Wenzel, & Brand, 2006; Schmidt et al., 2013). In the adult zebrafish telencephalon, Notch signaling is thought to keep cells quiescent (Type I cells) (Chapouton et al., 2011; Rothenaigner et al., 2011). Removal of Notch signaling leads to progression through activated (Type II cells) and transit amplifying (Type III) stages, followed by differentiation into distinct neural and glial populations, with final cell fate influenced by a combinatorial influence of multiple signaling systems (Chapouton et al., 2010). Work in the zebrafish has uncovered a key role for Wnt signaling in hypothalamic neurogenesis, with Wnt signaling being required for the formation of distinct neuronal subtypes and disruptions in Wnt-mediated hypothalamic proliferation leading to reduced growth and behavioral effects (Duncan et al., 2016; X. Wang et al., 2012; Xie & Dorsky, 2017). While Hh signaling has been shown to be a key component of hypothalamic-pituitary axis development during embryogenesis (Bergeron, Tyurina, Miller, Bagas, & Karlstrom, 2011; Blackshaw et al., 2010; Guner, Ozacar, Thomas, & Karlstrom, 2008; Kondoh et al., 2000; Muthu, Eachus, Ellis, Brown, & Placzek, 2016), its role in the post-embryonic and adult hypothalamus has not been documented.

Here we show that Hh signaling is necessary and sufficient to regulate hypothalamic neural precursor proliferation throughout the life cycle, with Hh-responsive cells being a previously undescribed population of proliferative radial glia similar to transit amplifying (type III) precursors. Using transgenic reporter lines we show remarkable heterogeneity in cell signaling responses among radial glia in the ventricular zone that appears to result in part from sequential responses of neural precursors to different signaling molecules. We show that Hh-responsive cells are multipotent neural precursors that give rise to dopaminergic, GABAergic, and serotonergic neurons. Together, these studies reveal a role for Hh signaling in larval and adult hypothalamic neurogenesis and begin to uncover heterogeneity among morphologically similar radial glial cells. Thus, our work may help explain how a wide range of neuroendocrine and neural populations can be regulated in the adult brain to allow for proper hypothalamic function and HP-axis regulation.

## Results

### Hedgehog responsive cells in the adult hypothalamus are proliferative neural precursors

We first took advantage of a suite of transgenic reporter lines to characterize cells that are actively engaged in Hh signaling in the adult brain (Fig. 1). Cells receiving Hh signaling (Hh responsive (Hh-R) cells) in the brain were visualized using transgenic reporter lines that express either GFP or mCherry under the control of the regulatory region of the *ptch2* gene, a transcriptional target of Hh signaling (Shen et al., 2013), while Sonic Hh (Shh)-producing cells were visualized using the *Tg(shha:GFP)* transgenic line (Neumann & Nuesslein-Volhard, 2000) (Fig. 1). Using modified CLARITY tissue clearing protocols (Isogai, Richardson, Dulac, & Bergan, 2017) and light sheet microscopy we obtained whole-brain images of adult *Tg(GBS-ptch2:NLS-mCherry;shha:GFP)* double transgenic fish, revealing that Hh-responsive (Hh-R) cells are adjacent to Shh producing cells in the posterior (PR) and lateral (LR) recesses of the hypothalamic ventricle (Fig. 1B). *in situ* hybridization confirmed that the *ptch2* reporter lines accurately report Hh signaling in the larval and adult brain, with *shh*, *ptch2* mRNA and the Gli transcription factors being expressed in ventricular regions (Supplemental Fig. 1). BrdU labeling of sagittal tissue sections revealed that Shh-producing cells are proliferative, have radial glial morphology, and that Shh-producing cells of the LR extend processes toward the PR (Fig. 1C). Similarly, Hh-responsive cells have radial morphology with processes that extend to the hypothalamic ventricle (Figs. 1D-H). Approximately 12% of these Hh-responsive cells in the adult PR are proliferative as revealed by expression of the Proliferative Cell Nuclear Antigen (PCNA) as well as BrdU incorporation (Fig. 1D, Fig. 3). A subset of Hh-R responsive cells of the posterior recess express the glial markers s100ß (Fig. 1E) and GFAP (Fig. 1F), while all appear to express the neural progenitor transcription factor Sox2 (Fig. 1G), consistent with neural progenitor identity. These cells give rise to neurons as shown by labeling with an antibody to the neuronal protein HuC (Park et al., 2000) (Fig. 1H). Together, these data show that a subset of radial glial cells in hypothalamic ventricular zones are responsive to Hh signaling in adults, and that these cells are both proliferative and neurogenic. These cells thus represent a previously undescribed population of neural precursors in the developing and mature vertebrate hypothalamus.

**Figure 1.**
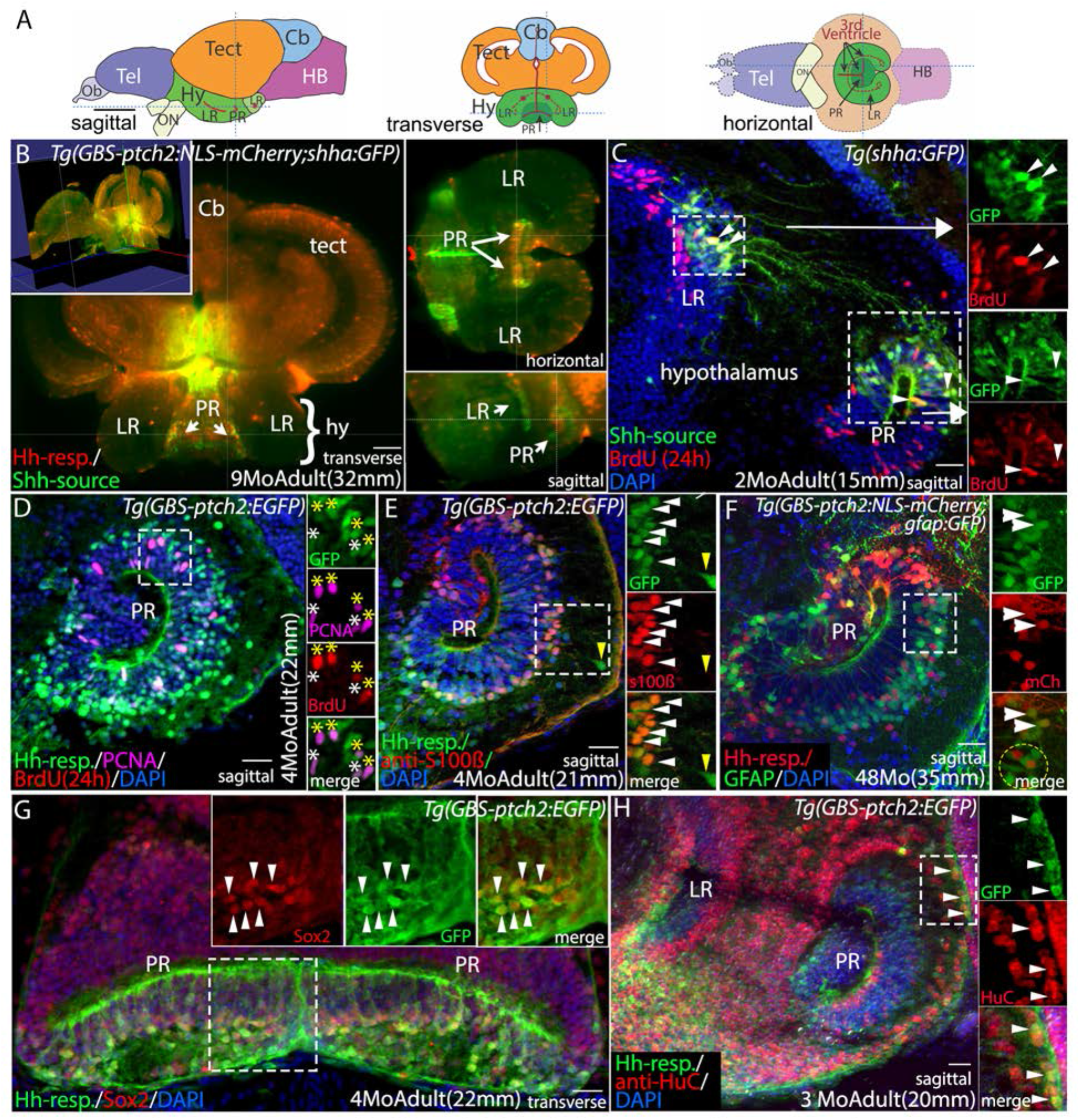
Shh-producing and Hh-responsive radial glial cells in the adult hypothalamus are proliferative neural precursors. **(A)** Schematic sagittal, transverse, and horizontal sections through the adult zebrafish brain. **(B)** Optical sections from a whole-brain light-sheet confocal image of a *Tg(GBS-ptch2:NLS-mCherry;shha:GFP)* double transgenic adult showing Hh-responsive cells (Hh-resp., red) in relation to Shh-producing cells (Shh-source, green) in the lateral recess (LR) and posterior recess (PR) of the adult hypothalamic (3rd) ventricle. Shh-expressing cells are adjacent to Hh-responsive cells in the ventricular zone of both recesses. Yellow lines in each panel indicate the plane of section in the other panels. (**C)** Sagittal tissue section showing Shh-producing cells (green) and BrdU labeled proliferative cells (red, 24-hour exposure) in the adult hypothalamus. Shh-producing cells are located primarily in the dorsal portions of both the lateral and posterior recesses. Shh-expressing cells in the LR send projections toward to the dorsal region of the PR. Panels at right show separated fluorescent channels from the boxed regions, with examples of co-labeled cells in both ventricular regions indicated by arrowheads. **(D)** Sagittal tissue section through the PR of a *Tg(GBS-ptch2:EGFP)* transgenic adult showing Hh-responsive cells and proliferative cells labeled with BrdU (red, 24h-hour treatment) and the anti-Proliferating Cell Nuclear Antigen (PCNA) antibody (magenta). Panels at right show separated channels from the boxed region, with 4 PCNA^+^/BrdU^+^ labeled Hh-responsive cells indicated by yellow asterisks and 2 PCNA^+^/BrdU^-^ cells indicated by white asterisks. **(E)** Sagittal tissue section through the PR of a *Tg(GBS-ptch2:EGFP)* transgenic adult labeled with the anti-s100ß antibody to show radial glia. Most but not all Hh-responsive cells are s100ß positive (white arrowheads). A small percentage of GFP labeled cells that are more distant from the ventricle are s100ß negative (yellow arrowhead), suggesting these cells have differentiated but retain GFP fluorescence from previous GBS-*ptch2:EGFP* transgene expression. **(F)** Sagittal tissue section through the PR of a *Tg(GBS-ptch2:NLS-mCherry;gfap:GFP)* double transgenic adult. About 5% (8 cells of 157 counted) of Hh-responsive cells in the adult hypothalamus express GFAP (white arrowheads), while the majority of Hh-responsive cells are GFAP negative (yellow circle). **(G)** Transverse tissue section through the PR of the hypothalamus of a *Tg(GBS-ptch2:EGFP)* adult labeled with the Sox2 antibody that labels neurogenic cells. Most or all Hh-responsive cells express the Sox2 protein. Panels at right show separated channels from the boxed region with arrowheads marking co-labeled cells. **(H)** Sagittal tissue section through the PR of a *Tg(GBS-ptch2:EGFP)* adult labeled with an antibody to the neuronal marker HuC/D (now called Elavl3). Double labeling of cells far from the ventricle indicates that Hh-responsive cells (green) can give rise to HuC/D expressing neurons (red). Panels at right show separated channels from the boxed region with co-labeled cells (arrowheads). Cb; cerebellum, hy; hypothalamus, LR: lateral recess of the hypothalamic (3^rd^) ventricle, PR; posterior recess of the hypothalamic (3^rd^) ventricle, tect; tectum. Scale bars: A, 1mm; B, 50µm; C-H, 20µm.

### Hedgehog signaling positively regulates hypothalamic neural precursor proliferation

To determine if Hh signaling affects adult neurogenesis we injected adult fish with the Hh inhibitor cyclopamine (CyA) (Incardona, Gaffield, Kapur, & Roelink, 1998) and assayed cell proliferation using PCNA antibody labeling and BrdU incorporation. These cyclopamine treatments dramatically reduced *ptch2* transcription as assayed by *in situ* hybridization (Supplemental Fig. 1), however green fluorescence was still visible in Hh-responsive cells after 24 hours of CyA treatment due to the persistence of the GFP protein (Fig. 2A,B). These IP injections of 80µM CyA reduced proliferation throughout the hypothalamus 24 hours later, with quantification of the posterior recess revealing an approximately 50% reduction in both BrdU and PCNA labeled cells (Fig. 2C).

**Figure 2.**
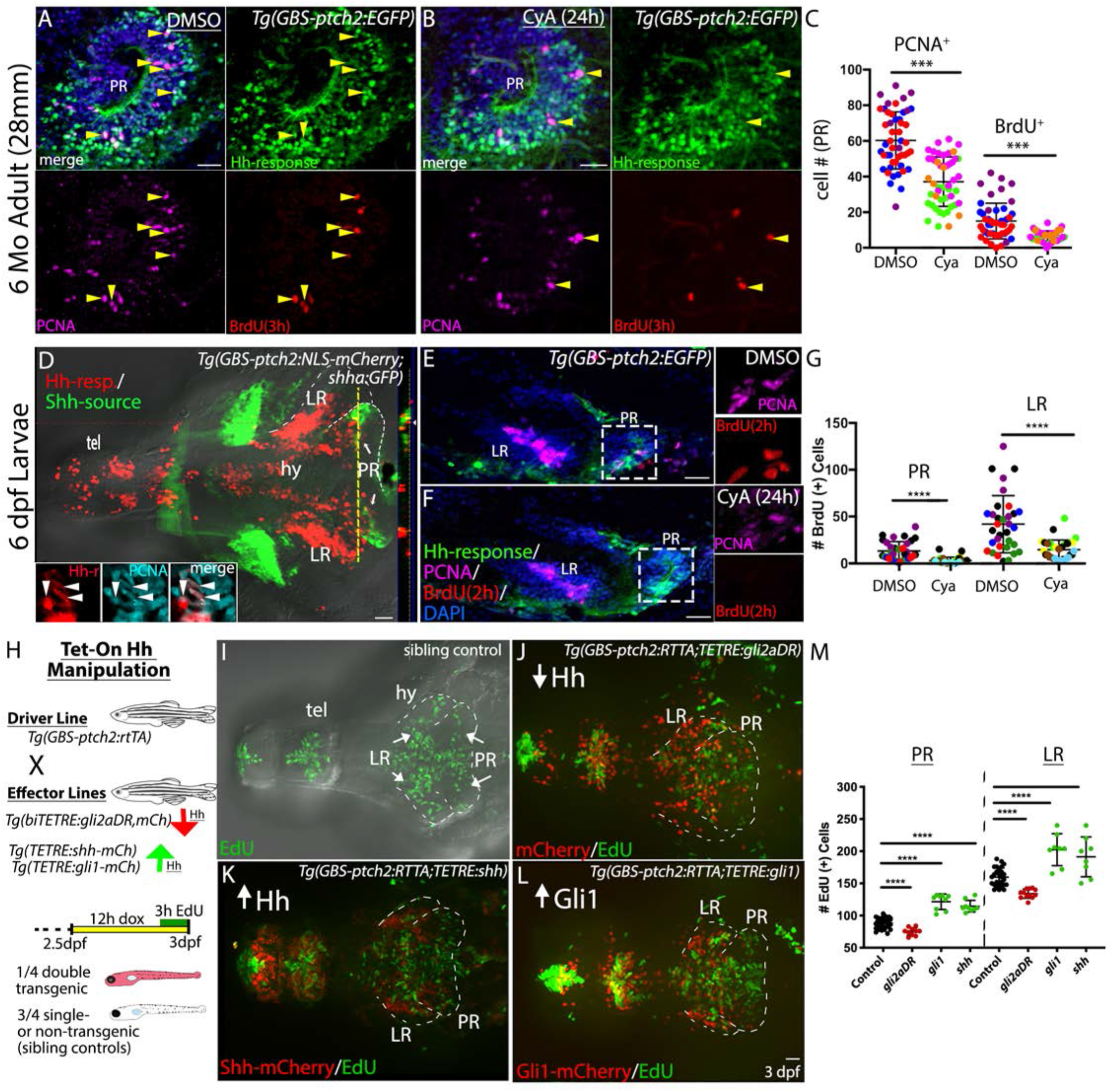
Hh signaling positively regulates proliferation in the postembryonic hypothalamus. **(A)** Sagittal section through the posterior recess of a DMSO (control) injected *Tg(GBS-ptch2:EGFP)* transgenic adult brain showing anti-PCNA (magenta) and BrdU (red, 3-hour exposure) labeled proliferative cells. Arrowheads mark triple labeled, proliferative, Hh-responsive cells. **(B)** Sagittal3-houron through the posterior recess of a *Tg(GBS-ptch2:EGFP)* transgenic adult brain 24 hours after injection of the Hh inhibitor cyclopamine (CyA) showing anti-PCNA (magenta) and BrdU (red) labeled proliferative cells. Arrowheads mark triple labeled cells. **(C)** Quantification of BrdU and PCNA labeled cells in control and CyA injected adults. Each dot represents the number of cells counted in a single tissue section, with each color representing a different adult fish (n=3 fish, 10-11 sections per fish). **(D)** Ventral view of a dissected *Tg(GBS-ptch2:NLS-mCherry;shha:GFP)* double transgenic larval brain showing Hh-responsive (red) and Shh-producing (green) cells. The hypothalamic lobes surrounding the posterior and lateral recesses of the third ventricle are outlined on one side of the brain. Cut view at right shows optical Z-section through the PR at the position of the yellow dotted line. Insets show PCNA labeled (cyan) Hh-responsive cells (red) in the PR of another larval brain, with examples of double labeled cells marked by arrowheads. **(E, F)** Representative sagittal tissue sections through the larval brains of DMSO (**E**) and cyclopamine (**F**) treated *Tg(GBS-ptch2:EGFP)* labeled to show BrdU (2h treatment) and PCNA labeled proliferative cells. Cyclopamine treatment led to a reduction in PCNA and BrdU-labeled cells in the PR, as well as throughout the brain (not shown). Panels at right show separated channels from the boxed region. **(G)** Quantification of BrdU and PCNA labeled cells in the PR and Lateral Recess (LR) of DMSO and cyclopamine treated larvae. Each color represents a different larval fish (n=5 controls, n=6 CyA treated) and each dot represents cells in a single tissue section (5-9 sections per fish). **(H)** Schematic showing the Tet-On transgenic system used to manipulate Hh signaling. The *Tg(GBS-ptch2:RTTA)* line drives expression of the RTTA transcriptional activator in Hh-responsive cells, with different effector transgenes allowing up- and down-regulation of Hh signaling upon the addition of doxycycline, which is required for RTTA function. Activation of the 2-part transgenic expression system is indicated by mCherry fluorescence in larvae, as shown in the diagram of the experimental timeline. **(I-L)** Ventral views of EdU labeled (3-hour exposure) proliferative cells in the larval hypothalamus following manipulation of Hh signaling levels using the Tet-On system. **(I)** EdU labeled proliferative cells (green) in a single-transgenic sibling (control), identified based on the lack of red fluorescence. **(J)** EdU labeled proliferative cells (green) visualized 12 hours after activation of a dominant repressor form of the Gli2 transcription factor (Gli2DR) in Hh-responsive cells (red). **(K)** EdU labeled proliferative cells (green) visualized following activation of the Shh effector transgene (*Tg(TETRE:shha-mCherry*)) in Hh-responsive cells *(Tg(GBS-ptch2:RTTA)* driver). **(L)** EdU labeled proliferative cells 12 hours after activation of the Gli1 transcription factor in Hh-responsive cells (red). **(M)** Quantification of cell proliferation in the PR and LR of the larval hypothalamus following Hh-manipulation using the Tet-On system. Each dot represents an individual fish. Up-regulation of Hh/Gli signaling via the *shh*a and *gli1* transgenes led to increased proliferation, while down-regulation of Hh signaling via the *gli2DR* transgene reduced proliferation. *** p<0.001. **** p<0.0001. Scale bars: 20µm.

We next turned to larval stages to facilitate gain- and loss- of function analyses of Hh function in the hypothalamus. As in adults, Shh-producing and Hh-responsive cells were distributed throughout the hypothalamic ventricle zones (Fig. 2D), with a subset being proliferative as revealed by PCNA labeling or BrdU incorporation (Fig. 2D, inset). CyA-mediated inhibition of Hh signaling at larval stages (6 dpf) reduced proliferation throughout the brain, with proliferation rates dropping by approximately 75% in both the posterior and lateral recesses of the hypothalamus (Fig. 2E-G). We saw no evidence of increased cell death in the hypothalamus of these CyA treated larvae, as assayed using an activated-Caspase3 antibody, although we did observe an intriguing increase of cell death in the dorsal tectum following CyA treatments (Supp. Fig. 2).

To verify the specificity of this effect we next used the two-part Tet-On system in zebrafish (Campbell, Willoughby, & Jensen, 2012), providing spatiotemporal control of Hh signaling levels (Fig 2). We previously demonstrated that expression of a truncated Gli2a transcription factor (Gli2aDR) dominantly represses Hh signaling at the transcriptional level, while expression of Shh or the full-length Gli1 transcription factor activates Hh signaling (Karlstrom et al., 2003; Shen et al., 2013). Expression of the Gli2aDR protein in Hh-responsive cells (*Tg(GBS-ptch2:RTTA-HA*) driver line) at 3 dpf resulted in a 25% reduction in hypothalamic proliferation within 12 hours of transgene activation (Fig. 2I,J). Similarly, activation of the Gli2a dominant repressor gene at 5 dpf using the heat-shock inducible system (Shen et al., 2013) reduced proliferation by approximately 50% (data not shown). Consistent with a positive role for Hh in regulating hypothalamic cell proliferation, expression of either Shha or the Gli1 transcriptional activator in Hh responsive cells starting at 3 dpf increased proliferation levels by approximately 20-30% within 12 hours of gene activation (Fig. 2K-M). Similar results were observed at 22 dpf, representing a late larval stage of development (data not shown). Together, these data indicate that Hh/Gli signaling is necessary and sufficient for normal proliferation rates in the brain at larval stages and that Hh continues to positively regulate cell proliferation in the adult hypothalamus.

### Hh-responsive cells are more highly proliferative than slow-cycling nestin-expressing radial glia

Nestin expressing radial glia have been identified as neurogenic cells in the zebrafish telencephalon (Chapouton, Jagasia, & Bally-Cuif, 2007; Ganz, Kaslin, Hochmann, Freudenreich, & Brand, 2010; Kaslin et al., 2009), with Nestin expression now being thought to be a hallmark of the transition from quiescent to activated neural stem cells in both mammals and teleosts (Chaker et al., 2016; Obernier & Alvarez-Buylla, 2019; Y. Z. Wang, Plane, Jiang, Zhou, & Deng, 2011). Nestin-expressing radial glia are also present in the hypothalamic ventricles, and these cells are largely PCNA negative (Fig. 3A), consistent with a quiescent or slow-cycling state. A higher percentage of Hh-responsive cells in the PR are PCNA positive (Fig. 3B, see quantification in 3F), with this PCNA labeling more prevalent toward the apical surface of the ventricular zone (Fig. 3B insets). This apical distribution of PCNA labeling is consistent with G1/S/G2 of the cell cycle occurring distant from the ventricle and M phase occurring near the ventricle, as has been shown in to occur during neurogenesis in the embryo (Alexandre, Reugels, Barker, Blanc, & Clarke, 2010; Leung, Klopper, Grill, Harris, & Norden, 2011). The fact that Hh-responsive cells and nestin positive cells have different cellular distributions and proliferative profiles (Fig 3 A,B) indicates that these cells are largely distinct. The persistence of GFP (X. Wang et al., 2012) and mCherry fluorescent proteins in the *TgBAC(nes:EGFP)* and *Tg(GBS-ptch2:NLS-mCherry)* lines allowed us to test the idea that these two transgenes might be sequentially expressed in the same cells as they progress through the stem cell activation-proliferation-differentiation pathway (Obernier & Alvarez-Buylla, 2019). We found that approximately 17% of mCherry expressing (i.e. Hh-responsive) cells in the periphery of the ventricular zone also contained GFP, indicating that these cells express, or previously expressed, the *nes:EGFP* transgene (Fig. 3C).

**Figure 3.**
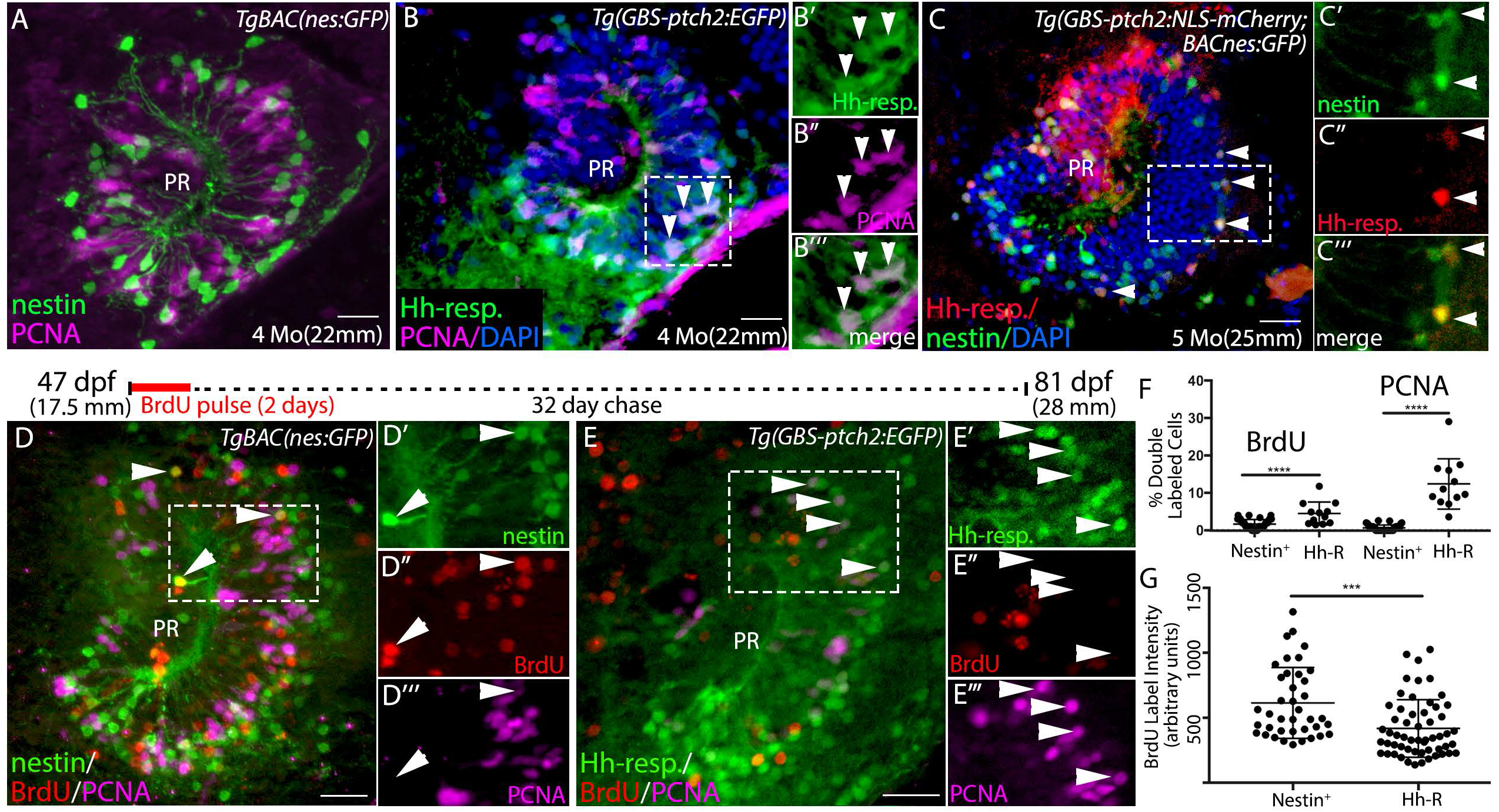
Hh-responsive progenitors are highly proliferative relative to slow-cycling nestin-expressing glia. **(A)** Sagittal tissue section through the PR of a *TgBAC(nes:EGFP)* transgenic adult brain showing PCNA antibody labeled proliferative cells. Nestin expressing cells are predominantly PCNA negative. **(B)** Sagittal tissue section through the PR of a *Tg(ptch2:EGFP)* transgenic adult brain showing PCNA antibody labeled proliferative cells. Arrowheads indicate examples of PCNA labeled (proliferative) Hh-responsive cells. Panels at right show separated channels from the boxed region**. (C)** Hh-responsive cells (red) are largely distinct from Nestin expressing cells (green) in the posterior recess (PR) of the adult hypothalamus, as visualized in *Tg(GBS-ptch2:NLS-mCherry;nes:EGFP)* double transgenic fish. However, 17% of cells (74 cells of 442 total) contained both GFP and mCherry (arrowheads, n=9 tissue sections from 3 double transgenic fish) with GFP fluorescence substantially lower in double-labeled cells. Panels at right show separated channels from the boxed region. **(D,E)** BrdU pulse-chase experiment. Schematic timeline above panels shows timing of pulse and chase, with 47 dpf adult *TgBAC(nes:EGFP)* or *Tg(GBS-ptch2:EGFP)* fish being exposed to 10 µm BrdU in fish water for 2 days. 32 days later fish were sacrificed and tissue sections were labeled using anti-BrdU (red) and anti-PCNA (magenta) antibodies. (**D)** Representative sagittal section through the posterior recess of a *TgBAC(nes:EGFP)* adult, insets show single channel data for the boxed region. A small number of Nestin-expressing cells in the posterior recess retained the BrdU label after one month. These cells did not express PCNA (arrowheads), indicating they were not in G1/S/G2 of the cell cycle at the time of fixation (n=3 fish, 13-16 tissue sections per fish). (**E)** Representative sagittal section through the posterior recess of a *Tg(GBS-ptch2:EGFP)* adult (n=2 fish, 13-16 sections per fish), insets show single channel data for the boxed region. Most Hh-responsive cells failed to retain the BrdU label after one month and many of these cells expressed PCNA (arrowheads), indicating active cell cycling at the time of fixation. **(F)** Graph showing the percentage of Nestin-expressing or Hh-responsive cells that co-labeled with the BrdU or PCNA antibodies. **(G)** Quantification of BrdU label intensity in Nestin expressing and Hh-responsive cells showing BrdU labeling intensity was significantly lower in Hh-responsive cells compared to nestin expressing cells. *** p<0.001. **** p<0.0001. All panels show 0.5 µm single optical sections of 20µm tissue sections. Scale bars: 20µm.

To experimentally determine the relative proliferation levels of nestin-expressing and Hh-responsive radial glia in the hypothalamus we performed BrdU pulse-chase experiments on *nestin:GFP* and *ptch2:EGFP* transgenic fish (Fig. 3). *TgBAC(nes:EGFP)* (Ganz et al., 2010) or *Tg(GBS-ptch2:EGFP)* (Shen et al., 2013) adults were treated with 10 mM BrdU for two days at 47 dpf and examined 32 days later to identify cells that retained the BrdU label, as well as cells that were proliferative at the time of fixation, as determined using the anti-PCNA antibody (Fig. 3 D-G). Consistent with low proliferation rates associated with activated stem cells, adult *nestin*-expressing cells in the PR very rarely (21/3853 cells counted, <1% of cells) expressed the proliferative marker PCNA, with approximately 2% of these cells retaining the BrdU label over the 1-month chase period, indicating they had gone through S-phase of the cell cycle one month earlier (Fig. 3D,F). Again, over 10% of Hh-responsive cells in the adult PR were found to be in G1/S/G2 of the cell cycle at the time of fixation based on PCNA expression, and approximately 5% of Hh-responsive cells had retained some BrdU label following the chase period, indicating they had progressed through S-phase of the cell cycle 1 month earlier. Fluorescent intensity of BrdU labeling in these label-retaining Hh-responsive cells was significantly lower than that in the BrdU+/*nestin*+ population (Fig. 3G), suggesting the Hh-responsive cells had undergone cell division(s) subsequent to the BrdU pulse period, thus reducing amount of the BrdU label in these cells. Together, these results support the hypothesis that *nestin*-expressing radial glia of the posterior recess represent a relatively slow cycling population while the Hh-responsive population represents a more rapidly cycling (e.g. Type III or transit amplifying stem cells) progenitor cell population.

### Hh, Wnt, and Notch signaling in hypothalamic progenitors

To explore potential interactions between Hh and other signaling pathways that control neurogenesis in the hypothalamus, we next determined the spatial relationship between Hh, Wnt and Notch in the ventricular zone. Examination of *Tg(7xTCF-Xla.Sia:GFP;GBS-ptch2:NLS-mCherry)* double transgenic adults revealed that the Wnt and Hh signaling systems are active in overlapping but distinct regions of PR, with Wnt-responsive cells being positioned primarily on the dorsal side of the ventricular zone and Hh-responsive cells being distributed throughout the ventricular zone (Fig. 4A). Close examination double transgenic adults revealed that a small subset of cells expressed both GFP and mCherry, indicating either simultaneous or sequential (given the persistence of the mCherry and GFP proteins) activation of Wnt and Hh signaling.

**Figure 4.**
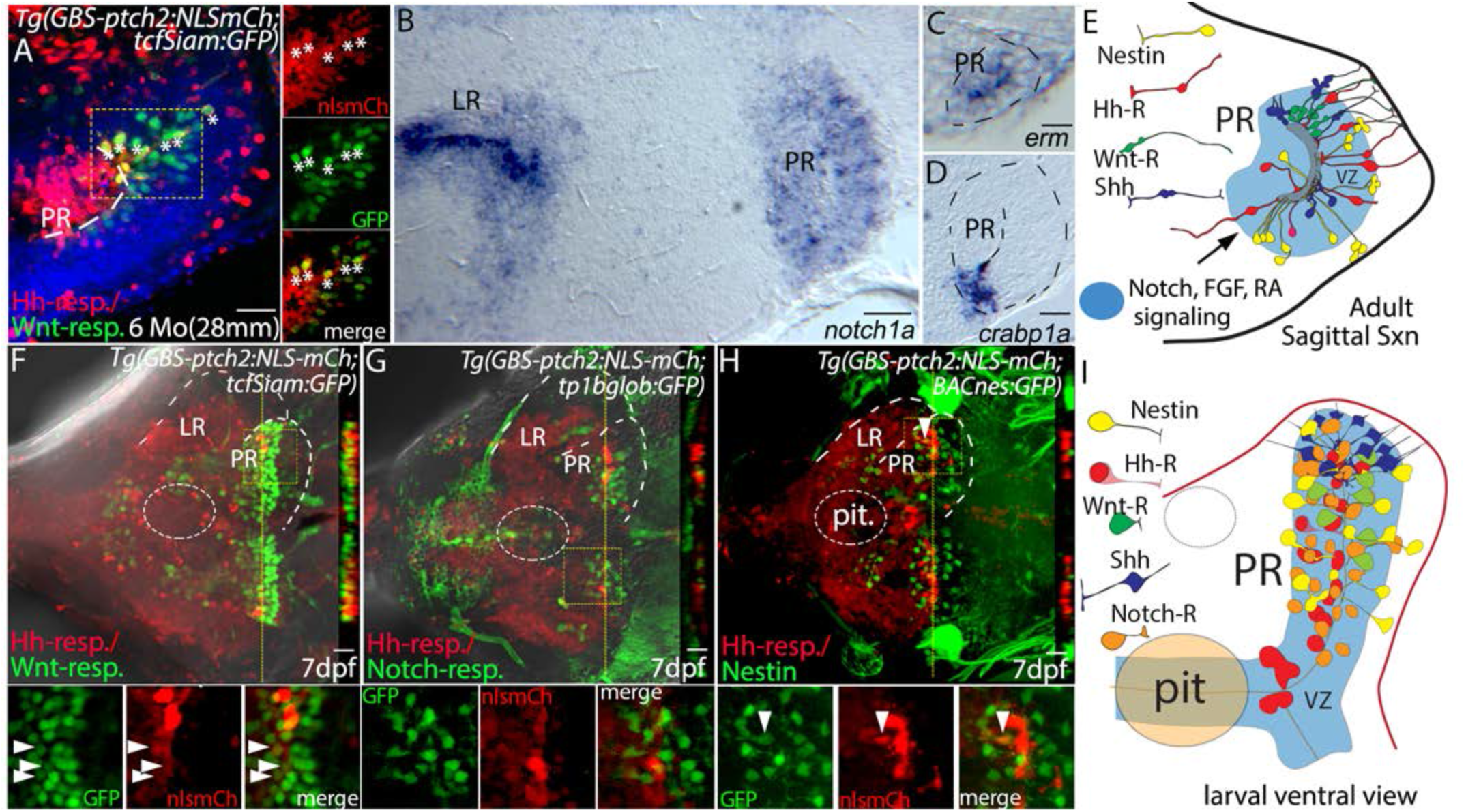
Hh, Wnt, Notch, FGF, and Retinoic Acid signaling in a complex hypothalamic neurogenic niche. (A) Sagittal section through the PR of a *Tg(GBS-ptch2:NLS-mCherry;TCFSiam:GFP)* double transgenic adult. Hh-responsive cells (red) of the PR are largely distinct from Wnt-responsive cells (green), however a subset of cells in the dorsal PR contains both GFP and mCherry (asterisks). Panels at right show separated channels from the boxed region. (B) Sagittal section through the hypothalamus showing *notch1a* expression in the LR and PR, as visualized by in situ hybridization. (C,D) Sagittal section through the PR showing expression of the FGF target gene *erm* (Raible & Brand, 2001) and the retinoic acid binding protein gene *crabp1a* ((R. Z. Liu et al., 2005). (E) Schematic of cell-cell signaling systems of the PR, including four distinct radial glial types that are defined by reporter gene expression (see Figs. 1-3 for data used to draw Shh-expressing and nestin expressing cells). (F-H) Ventral views of the 7 dpf hypothalamus of double transgenic larvae. Dotted lines outline the ventricular regions of the lateral (LR) and posterior (PR) recesses, and cut views at right show optical Z-sections through the PR at the position of the yellow dotted line. (F) Larval Hh-responsive cells of the PR are distinct from Wnt-responsive cells, as revealed in *Tg(GBS-ptch2:NLS-mCherry;TCFSiam:GFP)* double transgenic larva. (G) Hh-responsive and Notch-responsive cells of the PR are also distinct, as revealed in *Tg(GBS-ptch2:NLS-mCherry;tp1bglob:GFP)* double transgenic larva. (H) Hh-responsive cells of the PR are distinct from nestin expressing cells, as revealed in *Tg(GBS-ptch2:NLS-mCherry;nes:EGFP)* double transgenic larva. (I) Schematic ventral view of the larval hypothalamus showing cell-signaling pathways examined and four distinct radial glial types, as defined by gene expression in fluorescent reporter lines. pit; pituitary. Scale bars: 20µm.

Comparison of GFP expression between *TgBAC(nes:EGFP)* and *Tg(7xTCF-Xla.Sia:GFP)* adults revealed distinct patterns of expression with nestin+ cells distributed throughout the PR and wnt-R cells being primarily localized to the dorsal region (compare Fig. 3A to 4A), strongly suggesting that most Nestin expressing cells are not actively transducing Wnt signals. Given the data indicating that Notch signaling acts to keep telencephalic neural progenitors in a quiescent state (Chapouton et al., 2010), we next wanted to determine if Notch signaling might also be active in the zebrafish hypothalamus. *in situ* hybridization using the *notch1a* (Fig. 4B) as well as *notch1b* and *deltaB* (data not shown) probes revealed that Notch signaling is active in the LR and PR of the hypothalamic ventricle. The FGF target gene *erm* (Fig. 4C), as well as the gene encoding the retinoic acid binding protein Crabp1a (Fig. 4D), are also expressed in the adult PR, suggesting these signaling systems are actively involved in precursor regulation. A schematic summarizing the signaling environment of the adult posterior recess, as defined by these expression analyses, is shown in Fig. 4E.

We next examined whether the pattern observed in adults is be similar to that seen in the larval brain. Whole mount imaging revealed that Wnt-responsive and Hh-responsive progenitors are distributed throughout the PR, but these signaling systems appear to be largely acting on distinct cells, as indicated by the lack of co-expression in the Wnt and Hh-reporter lines (Fig. 4F). A very small number of double positive cells were identified in larvae (Fig. 4F, arrowheads), suggesting simultaneous or sequential activation of Wnt and Hh signaling in larvae similar to what we have observed in the adult. Notch-responsive cells, as seen in the *Tg(tp1bglob:GFP)* reporter line (Parsons et al., 2009), were present throughout the PR in a distributed pattern similar to that of Nestin expressing cells and this Notch-responsive population appears to be largely distinct from the Hh-responsive population (Fig. 4G). Finally, we observed that larval Nestin-expressing and Hh-responsive cell populations represent largely distinct precursor populations, but again with a small number of co-expressing cells (Fig. 4H arrowheads). Together, these results show that at least five embryonic cell-cell signaling systems, Hh, Wnt, FGF, RA, and Notch, persist throughout life and act on distinct radial glia populations in the posterior recess of the hypothalamus. The presence of a small number of co-labeled cells in double-transgenic larvae suggests that a subset of cells receive signals from multiple signaling systems, either simultaneously or sequentially. A schematic summarizing these data and the signaling environment of the larval posterior recess is shown in Fig. 4I.

### Hh-responsive cells are pluripotent neural progenitors that give rise to dopaminergic, GABAergic, and serotonergic neurons

To determine which neuronal subtypes are produced by Hh-responsive progenitors we crossed *Tg(GBS-ptch2:NLS-mCherry*) carriers to transgenic lines that express GFP in dopaminergic, all monoaminergic, GABAergic, or glutamatergic neurons. The persistence of fluorescent proteins for up to several days after termination of transgene transcription serves as a short-term lineage tracer and was used by others to demonstrate that Wnt-responsive cells in the PR give rise to both GABAergic and serotonergic neurons (X. Wang et al., 2012). Figure 5 A-C shows the presence of GFP/mCherry containing cells in double transgenic larvae as follows: *Tg(GBS-ptch2:NLS-mCherry;slc6a3:EGFP)* (labeling Dopaminergic cells, (Xi et al., 2011), *Tg(GBS-ptch2:NLS-mCherry;slc18a2:GFP)* (labeling monoaminergic cells (Wen et al., 2008), and *Tg(GBS-ptch2:NLS-mCherry; gad1b:GFP)* (labeling GABAergic cells (Satou et al., 2013). This indicates that Hh-responsive progenitors can give rise to dopaminergic neurons (Fig. 5A), possibly other monoaminergic neurons (Fig. 5B) and GABAergic neurons (Fig. 5C). Co-labeled dopaminergic cells were seen in both the lateral recess and posterior recess while co-labeled GABAergic cells were restricted to the lateral recess. In general, mCherry fluorescence was less intense in co-labeled cells, consistent with diminished quantities of mCherry protein that would be expected if these cells were no longer actively expressing the *GBS-ptch2:NLS-mCherry* transgene (i.e. were no longer Hh-responsive). We did not observe co-labeled cells in *Tg(GBS-ptch2:NLS-mCherry*;*slc17a6b:GFP)* double transgenic larvae, suggesting Hh-responsive cells may not give rise to glutamatergic neurons (data not shown). Finally, we used an anti-Serotonin antibody to label Serotonergic neurons in the *Tg(GBS-ptch2:NLS-mCherry*) line and found co-labeled cells in the PR, suggesting that Hh-responsive cells give rise to serotoninergic neurons (Fig. 5D). Together, these data reveal that Hh-responsive proliferative precursors are multipotent and can contribute to dopaminergic, serotonergic, and GABAergic populations in the hypothalamus.

**Figure 5.**
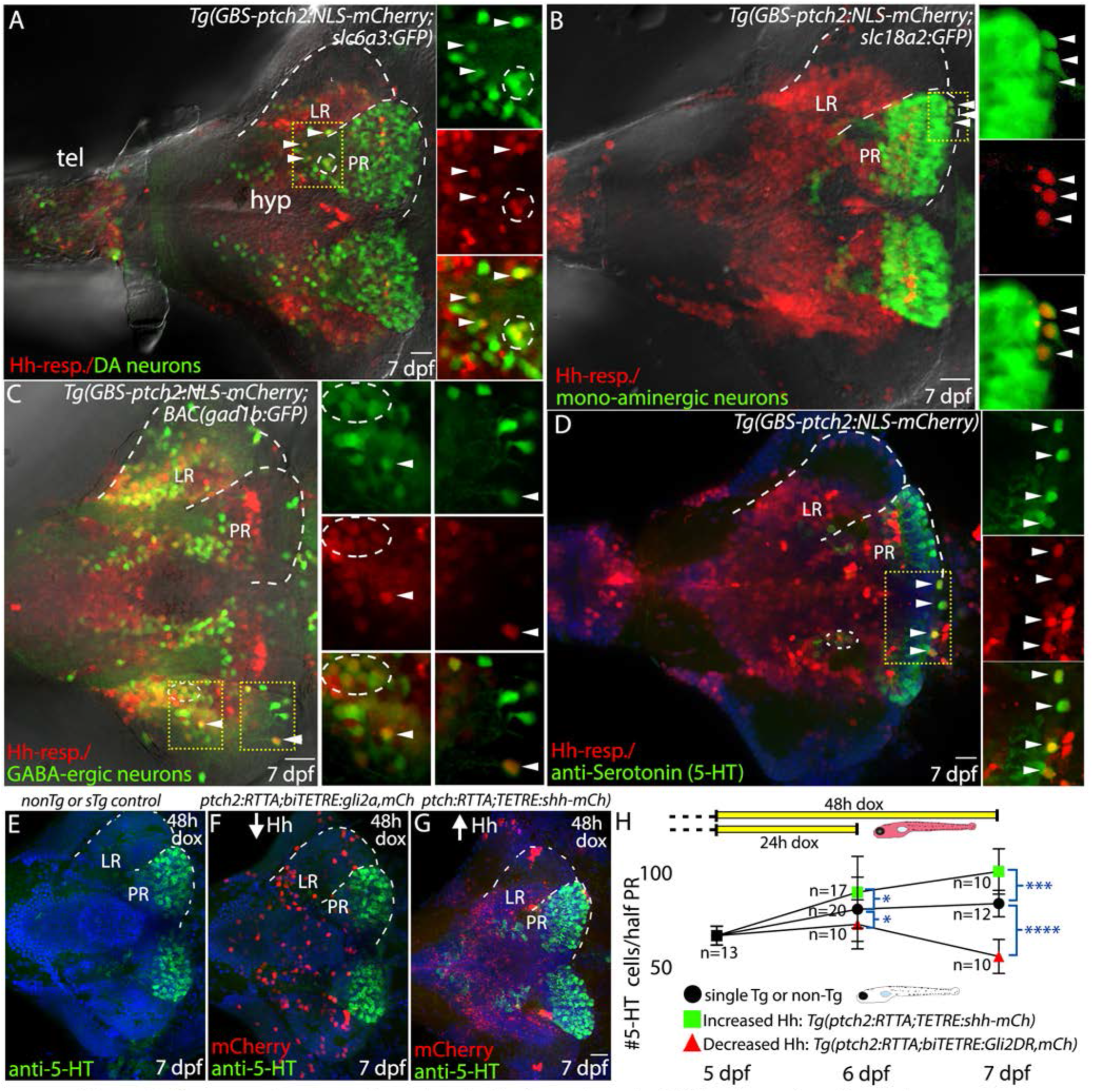
Hh-responsive progenitors of the hypothalamus give rise to dopaminergic, serotonergic, and GABAergic neurons. **(A)** Dopaminergic cells and Hh-responsive cells in the ventral brain, as visualized in a *Tg(slc6a3:EGFP,GBS-ptch2:NLS-mCherry)* double transgenic larva. A small subset of cells express both the GFP and the NLS-mCherry proteins (arrowheads, circle), suggesting Hh-responsive cells can give rise to dopaminergic neurons. (**B)** Monoaminergic neurons and Hh-responsive cells as visualized in a *Tg(slc18a2:GFP,GBS-ptch2:NLS-mCherry)* double transgenic larva. Again, a small subset of cells express both the GFP and the NLS-mCherry proteins (arrowheads), suggesting Hh-responsive cells can give rise to monoaminergic neurons. (**C**) GABAergic neurons and Hh-responsive cells as visualized in a *TgBAC(gad1b:GFP, GBS-ptch2:NLS-mCherry)* double transgenic larva. A small subset of cells express both the GFP and the NLS-mCherry proteins (arrowheads), suggesting Hh-responsive cells can give rise to GABAergic neurons. (**D**) Antibody labeling in a *Tg(GBS-ptch2:NLS-mCherry)* larval brain showing Serotonin (5-HT) expression in the ventral hypothalamus. The presence of double-labeled cells is consistent with Hh-responsive cells giving rise to serotonergic neurons. **(E-F)** Ventral views of anti-serotonin antibody labeled 7 dpf larval brains labeled cells following conditional manipulation of Hh signaling using the Tet-On transgenic system. **(E)** Representative single transgenic [*Tg(GBS-ptch2:RTTA-HA)* or *Tg(TETRE:shha-mCherry*)] sibling larva, identified by the lack of mCherry expression, showing number of serotonergic cells in the absence of effector transgene activation. **(F)** Representative *Tg(GBS-ptch2:RTTA,TETRE:shha-mCherry* double transgenic larva, identified by mCherry expression, showing increased numbers of serotonergic cells in the posterior recess following 2 day activation of the *shha-mCherry* transgene**. (F)** Representative *Tg(GBS-ptch2:RTTA,biTETRE:gli2aDR,nls-mCherry)* double transgenic larva, identified by mCherry expression, showing decreased numbers of serotonergic cells in the posterior recess following 2 day activation of the *gli2DR* transgene. **(H)** Graph showing serotonergic cell numbers, at 5 dpf, 6 dpf, and 7 dpf following 1 or 2 day activation of the Tet-On system in doxycycline (see diagram at top of graph). Error bars indicate standard deviation. Sample numbers for each experimental condition are shown on the graph, with significance determined using a one-way ANOVA. *;p<.05, ***;p<0.001, ****;p<0.0001. **(A-G)** Show ventral views of dissected brains from 7dpf larvae to show the hypothalamus. Dotted lines outline the lateral (LR) and posterior (PR) recesses. Small panels at right in A-D show single channel data for a single optical section in the boxed regions. hyp; hypothalamus, tel; telencephalon. Scale bars: 20µm.

### Hh signaling regulates serotonergic cell numbers in the hypothalamus

To determine if Hh signaling levels influence the generation of serotonergic cells in the hypothalamus we again employed our Tet-On Hh signaling transgenic lines. To facilitate antibody labeling, these experiments were terminated at 6 and 7 dpf. To repress Hh signaling levels in larval fish we crossed fish carrying the *GBS-ptch2:RTTA* driver transgene to fish carrying the *biTETRE:gli2aDR,NLS-mCherry* effector transgene and applied doxycycline to 5 dpf larvae for 24 or 48 hours to activate transgene expression. Activation of the Gli2DR transcriptional repressor prevented the addition of additional serotonergic neurons, with Gli2DR/mCherry expressing double transgenic larvae having 10% and 34% fewer serotonergic cells at 6 dpf and 7dpf, respectively, compared to non-mCherry expressing siblings at these same ages (single-transgenic or non-transgenic) (Fig. 5 E,F,H). To activate Hh signaling levels in larval fish we crossed fish carrying the *GBS-ptch2:RTTA* driver transgene to the *Tg(TETRE:shha-mCherry;myl7:EGFP)* individuals, again adding doxycycline to the resulting larvae at 5dpf to activate transgene expression. Consistent with a positive role for Hh in regulating differentiated neuronal populations in the hypothalamus, activation of Hh/Gli signaling for 24 or 48 hours led to a 11% and 21% increase, respectively, in serotonergic cell numbers in the PR of double transgenic larvae relative to non-mCherry expressing siblings (Fig. 5G,H). Taken together, these data indicate Hh signaling is necessary and sufficient to regulate the numbers of at least this neuronal cell type in the posterior recess of the hypothalamus.

## Discussion

### Hedgehog signaling regulates neural stem cell proliferation in the adult hypothalamus

Gli-mediated Hh signaling plays a major role in formation of the hypothalamic-pituitary axis during embryonic development across vertebrate species (Burbridge, Stewart, & Placzek, 2016; Corman, Bergendahl, & Epstein, 2018; Devine et al., 2009; Guner et al., 2008; Sbrogna, Barresi, & Karlstrom, 2003) and a role for Hh signaling in regulating adult neural stem cell proliferation in the dorsal regions of the mammalian brain is now well established (Ahn & Joyner, 2005; Ihrie et al., 2011; Machold et al., 2003). Here we demonstrate for the first time a role for Hh/Gli signaling in adult hypothalamic neurogenesis. Hh responsive radial glial cells in the ventricular zone of the larval and adult hypothalamus are relatively rapidly proliferating multipotent neural progenitors that give rise to dopaminergic, serotonergic, and GABAergic neurons. Finally, conditional manipulation of Hh signaling revealed that Hh signaling is necessary and sufficient for driving hypothalamic proliferation and that Hh signaling positively regulates the production of at least serotonergic neurons in the hypothalamus. Shh expression in DA neurons is required for DA neuron function/survival in mice, suggesting conserved Hh function in this cell type (Gonzalez-Reyes et al., 2012).

While approximately 10% of Hh responsive cells were found to be proliferative in adults, we also found a substantial number of Hh-responsive cells that retained a BrdU label for more than a month (Fig. 3E). This indicates substantial heterogeneity in the proliferative rates of Hh-responsive cells that could be due to dose-dependent responses to Hh ligands and/or cross regulation by as yet unidentified mechanisms. Given the local and distant sources of Shh in the PR region, Shh producing cells may act as key modulators of hypothalamic proliferation rates, both globally to support hypothalamic growth and tissue renewal, and regionally to regulate region specific growth rates and to control the generation of specific cell types needed for hypothalamic function. Indeed, loss of Hh signaling in larvae led to both reduced proliferation rates and the reduction in the size of the serotonergic population. These data suggest that regulation of Hh signaling levels could play a role in life-long hypothalamic plasticity and function.

### Hh-signaling and Hypothalamic Stem Cell Progression

We show that Nestin-expressing radial glia represent a relatively slow cycling progenitor population that retains BrdU label for one month, thus resembling quiescent/activated neural stem cells in more dorsal regions of the mammalian brain (Ming & Song, 2011) (Wang et al., 2011). The fact that a small percentage of nestin expressing cells were also PCNA positive is consistent with the nestin cells representing an early activated neural stem cell population (Daynac et al., 2016). In the zebrafish and mammalian telencephalon Notch signaling acts to keep stem cells in a quiescent state, with inhibition of Notch activating these cells as a first step in the stem cell proliferation pathway (Carlen et al., 2009). We show that Notch signaling is active in the hypothalamic ventricular zone throughout life and that Nestin-expressing and Notch-responsive cells are distributed throughout the posterior recess in a pattern that is distinct from the Hh-responsive population (Fig. 4). The relative numbers of cells, proliferation rates, and distribution of these cell types is consistent with a role for Notch in regulating early steps (e.g. activation) in stem cell progression (Chapouton et al., 2010), with Hh/Gli then acting further downstream to control proliferation rates and possibly differentiation (Alvarez-Medina, Le Dreau, Ros, & Marti, 2009; Corman et al., 2018; Locker et al., 2006; Palma et al., 2005).

We observed that a small percentage of nestin expressing cells were Hh responsive, with relative fluorescent protein intensity consistent with the Hh-response occurring in cells that previously expressed Nestin. Combined with the finding that Hh-responsive cells are more highly proliferative than their neighboring nestin-responsive cells, this suggests a model in which Hh/Gli signaling begins in the activated stem cell population and regulates subsequent proliferation rates in a Hh-responsive transit amplifying population, likely acting as a mitogen to regulate cell cycle progression. Hh signaling was recently shown to shorten G1 and S-G2/M portions of the cell cycle in the mammalian sub ventricular zone, results that were interpreted as showing a role in neural stem cell activation (Daynac et al., 2016). However, these mammalian data are also consistent with a role for Hh signaling in precursor amplification, similar to the role we show here in the hypothalamus.

### The hypothalamic posterior recess: a complex neural progenitor niche regulated by multiple signaling molecules

In mammals, Hh, Notch, and Wnt signaling systems have all been shown to remain active in the adult hypothalamus (Mirzadeh et al., 2017) with FGF-signaling linked to postnatal hypothalamic proliferation and affecting energy balance and appetite (Robins et al., 2013). We and others have now shown that Wnt, Hh, Notch, FGF, and RA signaling systems are all active in the hypothalamic ventricular zone of larval and adult zebrafish (this study, (Guner et al., 2008; Shearer et al., 2010; Topp et al., 2008; X. Wang, Lee, & Dorsky, 2009), indicating this region represents a complex and heterogeneous signaling environment similar to mammalian stem cell niches (Chaker et al., 2016). Our data demonstrating that Wnt, Hh, and Notch signaling act on distinct but overlapping populations of radial glia (see Fig. 4) reveals heterogeneity in progenitor populations and suggests distinct, possibly sequential functions in regulating activation and proliferation of precursors. Given the importance of adult neurogenesis in hypothalamic function, it will now be important to explore the relative roles of these signaling molecules in regulating neurogenesis.

Wnt signaling was previously shown to play a role in neurogenesis within the zebrafish hypothalamic ventricle {Wang, 2009 #2588;Wang, 2012 #4428}, with Wnt signaling affecting neuronal differentiation and possibly activation of quiescent stem cell populations, but with little evidence for Wnt signaling in controlling proliferation rates (Duncan et al., 2016). In other systems Hh and Wnt signaling pathways are known to act on proliferation in the same cells (Alvarez-Medina et al., 2009). While expression profiles suggest the Wnt and Hh/Gli signaling systems may act on distinct radial glia in the hypothalamus, our finding that a small number of cells are co-labeled in the Wnt and Hh reporter lines is consistent with sequential action of Wnt and Hh signaling, with Wnt acting on the early activation of quiescent cells, followed by Hh regulating the level of progenitor amplification prior to differentiation. The results presented here set the stage for detailed analyses of how these multiple signaling pathways combine to regulate stem cell activation, proliferation, and differentiation of distinct neural stem cell populations.

### Adult neurogenesis and hypothalamic function

A number of studies have now linked proliferation in the hypothalamus to distinct hypothalamic functions, including regulation of energy metabolism (Kokoeva et al., 2005; Kokoeva, Yin, & Flier, 2007), anxiety in zebrafish (Xie et al., 2017) and seasonal reproductive changes in sheep (Migaud et al., 2011). These changes in proliferation rates must be tightly regulated in response to season or changing metabolic needs. Sonic Hedgehog and other embryonic cell-cell signaling systems are positioned as possible mediators of short-term plasticity in neuronal populations that contributes to homeostasis. Since HP axis function relies on the regulated output of a large number neurosecretory cells, regulating cell numbers of distinct populations may be an important component of normal HP axis homeostasis. Consistently, hypothalamic pro-opiomelanocortin (POMC) populations were shown to change in response to altered temperature, impacting feeding (Jeong et al., 2018). Similarly, regulating dopaminergic neuron numbers could affect responses to osmotic challenges (N. A. Liu et al., 2006) while changes in serotonergic neuronal populations may be linked to stress responses (Xu et al., 2019). Our data add to a growing body of evidence that link embryonic cell-signaling systems to the regulation of adult neuronal populations needed for life-long homeostasis and metabolic health, with coordinated mitogenic activity among these multiple signaling systems potentially underlying adult hypothalamic plasticity critical for the vertebrate response to metabolic challenges.

## Materials and Methods

### Zebrafish strains and transgenic lines

Zebrafish were maintained as described previously (Kimmel et al., 1995) in compliance with the University of Massachusetts, Amherst, Institutional Animal Care and Use Committee protocols. Wild type lines used were TL and AB. Transgenic lines used are listed in table 1.

**Table 1:** Transgenic Lines Used in this study

| Transgenic Line | Reference |
| --- | --- |
| *Tg(GBS-ptch2:EGFP)<sup>umz231g</sup>* | (Shen et al., 2013) |
| *Tg(GBS-ptch2:NLS-EGFP)<sup>umz241g</sup>* | (Shen et al., 2013) |
| *Tg(GBS-ptch2:mCherry)<sup>umz261g</sup>* | (Shen et al., 2013) |
| *Tg(GBS-ptch2:NLS-mCherry)<sup>umz271g</sup>* | (Shen et al., 2013) |
| *Tg(hsp70l:shha-EGFP)<sup>umz301g</sup>* | (Shen et al., 2013) |
| *Tg(hsp70l:gli1-EGFP)<sup>umz311g</sup>* | (Shen et al., 2013) |
| *Tg(hsp70l:gli2aDR-EGFP)<sup>umz331g</sup>* | (Shen et al., 2013) |
| *Tg(GBS-ptch2:RTTA-HA)<sup>umz391g</sup>* | This paper |
| *Tg(actb2:RTTA-HA;myl7:EGFP)<sup>umz401g</sup>* | This paper |
| *Tg(TETRE:shha-mCherry;myl7:EGFP)<sup>umz411g</sup>* | This paper |
| *Tg(biTETRE:gli1,NLS-mCherry)<sup>umz421g</sup>* | This paper |
| *Tg(biTETRE:gli2aDR,NLS-mCherry)<sup>umz431g</sup>* | This paper |
| *Tg(-2.7shha:GFP)<sup>t101g</sup>* | (Neumann & Nüsslein-Volhard, 2000) |
| *TgBAC(nes:EGFP)<sup>tud1001g</sup>* | (Ganz et al., 2010) |
| *Tg(Tp1bglob:eGFP)<sup>um13</sup>* | (Parsons et al., 2009) |
| *Tg(gfap:GFP)<sup>mi2002</sup>* | (Bernardos & Raymond, 2006) |
| *Tg(slc6a3:EGFP) aka Tg(DAT:gfp)* | (Xi et al., 2011) |
| *Tg(7xTCF-Xla.Siam:GFP)<sup>ia4</sup>* | (Moro et al., 2012) |
| *Tg(slc18a2:GFP) or Vmat2:GFP* | (Wen et al., 2008) |
| *Tg(slc17a6b:DsRed)<sup>nns9</sup> aka Vglut:dsRed* | (Satou et al., 2013) |
| *TgBAC(gad1b:GFP)<sup>nns25</sup>* | (Satou et al., 2013) |

### Generation of Tet-On Hedgehog-pathway manipulation transgenic lines

Transgene constructs were constructed using Gateway cloning system. The destination and entry clones containing EGFP or mCherry were from the zebrafish Tol2kit systems (Kwan et al., 2007; Villefranc, Amigo, & Lawson, 2007). Entry clones for the Hh-pathway specific elements are described in (Shen et al., 2013) and entry clones with Tet-On system elements were generously provided by the Jensen lab (Campbell et al., 2012). Briefly, 25 pg of purified plasmid DNA was injected to one-to two-cell zebrafish embryos with 25 pg of transposase mRNA. Injected fish were raised to adulthood and out-crossed to wild-type individuals to identify potential founder fish. The transgenic fish containing the *myl7:EGFP* transgene were identified by GFP expression in heart; other lines were PCR genotyped using the primer sets shown in Table 2.

**Table 2:**
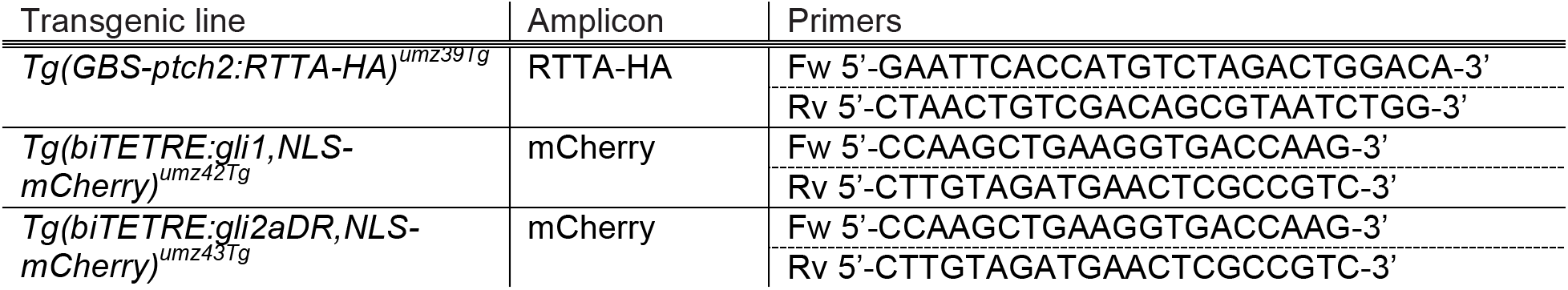
Genotyping Primers

### Tissue preparation, sectioning, imaging, and cell counts

Larvae and adults (heads only) zebrafish were anesthetized using tricaine methanesulfonate (Sigma-Aldrich), and fixed overnight in 4% paraformaldehyde (PFA) at 4 °C. For tissue sectioning, samples were washed three times in PBS/0.1% Tween (PBST), placed in embedding media (1.5%agar, 5%sucrose), incubated in 30% sucrose solution overnight at 4°C, and 20μm sections were cut using a Leica CM 1950 cryostat. Larval brains were dissected after fixation and mounted in 25% glycerol. Imaging was performed using a Zeiss Axioskop 2 Apotome or LSM 700 laser scanning confocal microscope. Images are single optical planes except where noted. For lightsheet imaging, adult brains were fixed overnight in the cranium, dissected, embedded in acrylamide gel and cleared as in (Isogai et al., 2017). Lightsheet images were collected as optical sectioned z-stacks using a Zeiss Lightsheet Z1. Cell numbers and fluorescent intensity were quantified manually using ImageJ software (NIH) on single optical sections. Cells were outlined and the mean fluorescent intensity for each cell was calculated by drawing a circle of defined area through the brightest plane containing the nucleus.

### Cyclopamine (CyA)

3-7 day old larvae were exposed to 75-100µM cyclopamine (Toronto Chemical) (Incardona et al., 1998) by adding 10 µl of 10 mM stock solution (CyA dissolved in DMSO) to 1 ml of embryo rearing medium (Westerfield, 2007) in 12 well tissue culture plates (∼10 larvae/well) at 28.5 °C. Control larvae were treated with equal volumes (10 µl) of DMSO. Adults were injected intraperitoneally with a concentration of 0.2mg/ml in a volume of 25 µl (10 mg/kg) CyA in DMSO (Reimer et al., 2009). Control animals were injected with DMSO.

### Conditional Gene activation using the Tet-On system

Progeny derived from adults carrying an RTTA-encoding driver transgene *(Tg(GBS-ptch2:RTTA-HA))* or *Tg(actb2:RTTA-HA;myl7:EGFP)* and a TRE-containing effector transgene *(Tg(TETRE:shha-mCherry; myl7:EGFP), Tg(biTETRE:gli1,NLS-mCherry) or Tg(biTETRE:gli2aDR, nls-mCherry)* were sorted by GFP expression in the heart (*myl7:EGFP* transgene containing lines). RTTA-mediated gene expression was activated in 3-4 dpf larvae by adding 50µg/mL (final concentration) doxycycline to system water (commonly used embryo media contained calcium and magnesium and can cause precipitation of doxycycline). Doxycycline-containing water was replaced every 12 hours. Double transgenic embryos were identified by the presence of mCherry fluorescence using a Leica fluorescent dissecting microscope.

### Bromodeoxyuridine (BrdU) and Ethynyldeoxyuridine (EdU) Labeling

3-7 day old larvae were bathed in 10mM BrdU or 3.3mM EdU for 1-3 hours, while adults were bathed for 2 days in 10 mM BrdU dissolved in 0.5% DMSO. Stocks solutions were 100mM and 33.3mM, respectively. Following fixation in 4% PFA BrdU was detected in tissue sections using the rat anti-BrdU (Abcam) or mouse anti-BrdU G3G4 (Developmental Studies Hybridoma Bank) antibodies at dilutions of 1:300 and 1:10, respectively. EdU was detected in fixed, dissected larval brains as described in the Click-It EdU Labeling kit (Invitrogen).

### Immunohistochemistry and in situ hybridization

Whole-mount larval immunohistochemistry was performed as in (Guner & Karlstrom, 2007) with a few changes. For anti-PCNA labeling dissected whole larval brains were incubated in 1x Histo-VT One (Nacalai Tesque) for 60 min at 65 °C as in (Than-Trong et al., 2018). For anti-serotonin labeling dissected brains were digested in 60ug/ml proteinase K for 30 minutes at room temperature before proceeding for whole mount immunohistochemistry. Immunohistochemistry on sectioned larval and adult tissue was performed as in (Barresi, Hutson, Chien, & Karlstrom, 2005). Primary antibodies used were: rabbit anti-serotonin (1:1000, #20080, ImmunostarInc, WI, USA), mouse anti-S100β (1:500, #S50430-2, Dako/Agilent, CA, USA), rabbit anti-BLBP (1:300 #ABN14, MilliporeSigma, MA, USA), mouse anti-Sox2 (1:200, #S1451 Sigma Aldrich, MA, USA), rabbit anti-PCNA (1:50, #FL-261, Santa Cruz, Texas, USA), mouse anti-PCNA (1:500 Sigma, #p8825, MilliporeSigma, MA, USA), mouse anti-GFP (1:100, #MA5-15256, ThermoFisher Scientific, MA, USA), rabbit anti-GFP (1:500, #TP401, Torrey Pines Origene Technologies, Inc, MD, USA or #A-11122 ThermoFisher Scientific, MA, USA for), mouse anti-HuC/D (1: 300, #A-21271 ThermoFisher Scientific, MA, USA), rabbit anti-GFAP (1:200 gift from Nona lab). Secondary antibodies raised in goat were used at a 1:1000 dilution. *in situ* hybridization was performed as described previously (Shen et al., 2013). Probes used were: *notch1a* (Raymond, Barthel, Bernardos, & Perkowski, 2006), *erm* (Raible & Brand, 2001)*, crabp1a (R. Z. Liu et al., 2005)*, *shha (Krauss, Concordet, & Ingham, 1993)*, *ptch2* (Concordet et al., 1996), and *gli1*, *gli2a*, and *gli3* (see (Devine et al., 2009).

### Statistical Analyses

Cell numbers are represented as mean ± SD (standard deviation). Statistical significance is defined as * p ≤ 0.05, ** p < 0.01, *** p < 0.001, **** p < 0.0001. Student’s *t*-test was used for pairwise comparisons between two groups. One-way ANOVA with Tukey’s multiple comparisons (Prism 6.0 GraphPad software) was used to determine statistical significance of multiple comparisons.

## Acknowledgements

We would like to thank the Karlstrom lab members for stimulating discussions. We are grateful to Dr. Joseph Bergan for making the light sheet imaging possible, to Dr. Tatjana Piotrowski and Dr. Laure Bally-Cuif for sharing EdU and PCNA labeling protocols, and to Drs. Nathan Lawson, Richard Dorksy and Florian Engert and the zebrafish community for sharing *transgenic* zebrafish lines. We are particularly thankful to Caroline Wee of the Engert lab for her help providing fish. We thank Dr. Desmond Ramirez for the kind gift of the anti-serotonin antibody.

## Author Contributions

I.M. and A.T.O. designed experiments, collected, analyzed, and interpreted the data, and contributed to the writing of the manuscript, with A.T.O establishing and I.M. completing the project. R.R.F and M.L. performed some experiments, and collected and interpreted data, M-S.C. and V.P. established and validated the Tet-On transgenic lines, A.L. and C.D contributed expression data. R.O.K guided the project, designed experiments, interpreted data, and wrote the manuscript.

---

**Supplemental Figure 1.**
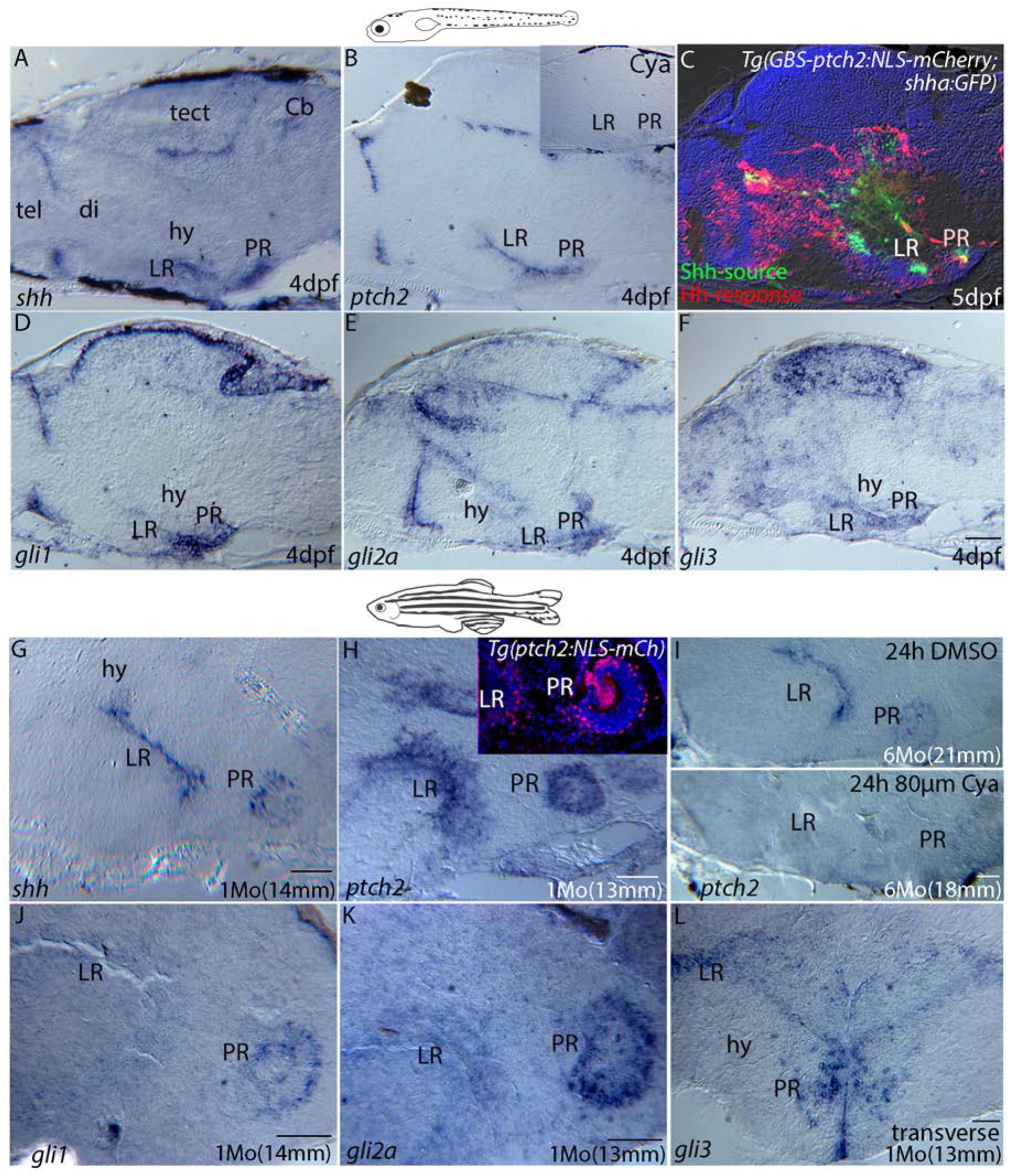
Hedgehog signaling pathway gene expression in the larval and adult hypothalamus. **(A-F)** Expression of Hh-signaling pathway genes in sectioned tissue from 4 dpf larvae. **(A)** *in situ* hybridization (ISH) showing that *shh* is highly expressed in the ventricular regions of the lateral recess and posterior recess of the hypothalamic ventricle, as well as in ventricular regions of the diencephalon/telencephalon border, cerebellum, and tectum. **(B)** The Hh-target gene *patched2* is similarly expressed in ventricular regions throughout the larval brain, with *ptch2* transcription being eliminated by treatment with cyclopamine **(B Inset). (C)** *shh* (green, Shh-source) and *ptch2* (red, Hh-response) expression in the larval brain as revealed by the *Tg(shha:GFP)* and *Tg(GBS-ptch2:NLS-mCherry)* reporter lines, seen here in a double transgenic larva. This midline section reveals Hh-response in the midline ventricular region**. (D-F)** ISH showing expression of the Hh-responsive transcription factors *gli1*, *gli2a*, and *gli3*, respectively. **(G-L)** *in situ* hybridization (ISH) on tissue sections showing expression of Hh signaling pathway genes in the adult hypothalamus. **(G)** *shh* expression is maintained in the lateral and posterior recesses of the adult hypothalamic ventricle. **(H)** Expression of the Hh-target gene *patched2* in the hypothalamic ventricular regions as revealed by ISH and compared to nuclear mCherry expression in cells of the LR and PR driven by the *ptch2* promoter construct in the *Tg(GBS-ptch2:NLS-mCherry)* transgenic line (**inset**). **(I)** ISH using a *ptch2* probe shows that cyclopamine treatment (bottom panel) of 6-month-old adult dramatically reduced *ptch2* gene expression in the lateral and posterior recesses compared to an age-matched DMSO control treated fish (top panel). **(J-L)** ISH showing expression of the Hh-responsive transcription factors *gli1*, *gli2a*, and *gli3*, respectively, in the hypothalamus. All panels show sagittal tissue sections, except L, which shows a transverse tissue section. cb; cerebellum, di; diencephalon, hy; hypothalamus, LR; lateral recess, PR; posterior recess, tect; tectum, tel; telencephalon. Scale bars: 50µm.

**Supplemental Figure 2.**
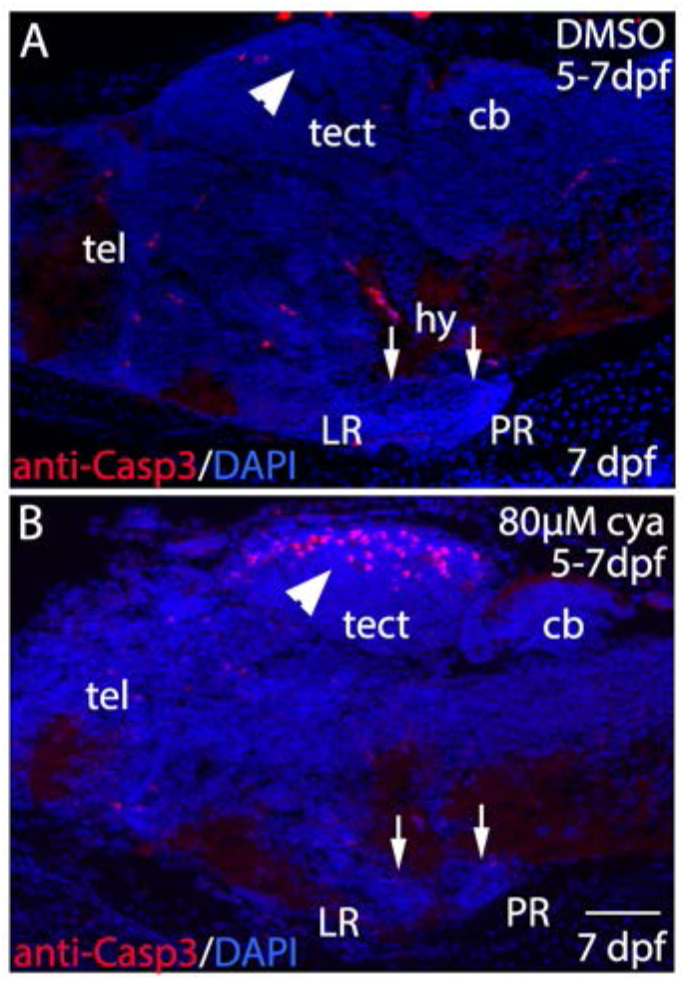
Cyclopamine treatment does not increase cell death in the hypothalamus. **(A)** Cell death in a DMSO treated control larva as revealed by anti-activated Caspase3 antibody labeling. **(B)** No cell death was seen in the hypothalamus (arrows) after two-day cyclopamine treatment, while increased cell death was seen in the tectum (arrowheads). cb; cerebellum, hy; hypothalamus, LR; lateral recess, PR; posterior recess, tect; tectum, tel; telencephalon. Scale bar: 50µm.

